# Human promoter analysis of the Programmed Axon Death genes *NMNAT2* and *SARM1*

**DOI:** 10.64898/2026.03.23.712947

**Authors:** Laura Carlton, Heba Morsy, Jonathan Gilley, Henry Houlden, Mary M. Reilly, Michael P. Coleman, Emma R. Wilson

## Abstract

SARM1 and NMNAT2 are two well described players in the Programmed Axon Death (PAxD) pathway. However, less is known about their transcriptional regulation, especially in humans, despite evidence that their expression levels influence axon vulnerability and thus modulation of expression presents a potential therapeutic target. Here, we used in-cell luciferase assays to functionally study the promoter regions of the human *NMNAT2* and *SARM1* genes. We find that human *NMNAT2* expression can be driven by cAMP, acting through one cAMP response element (CRE), compared to two in mice. Naturally occurring single-nucleotide variants exist within the CRE, some of which lower *NMNAT2* promoter activity by more than 50%. We also report an ultra-rare single nucleotide variant in the *NMNAT2* promoter in an ALS patient in Project MinE. This variant demonstrates pathogenic potential by lowering *NMNAT2* promoter activity in our assay. Project MinE also reveals a common *SARM1* promoter variant that significantly increases *SARM1* promoter activity in our assay. Thus, several single nucleotide changes in the *NMNAT2* and *SARM1* promoters modify transcription levels in the direction that would predict an increase in susceptibility to PAxD. These promoter variants refine our understanding of regulatory mechanisms affecting *NMNAT2* and *SARM1* expression and, together with previously reported coding variants for these genes, expand the catalogue of functionally relevant variants for future association studies in neurodegenerative diseases, including peripheral neuropathies and motor nerve disorders.

## Introduction

The Programmed Axon Death (PAxD) pathway appears to be a mechanistic convergence point for numerous inherited and acquired neurodegenerative diseases and a prospective therapeutic target (Coleman, 2022; Loreto & Neukomm, 2025). SARM1, an NAD degrading enzyme, is the central executioner of this pathway, which when activated causes cell death. SARM1 is maintained in an autoinhibited state by the binding of its substrate, NAD, to an allosteric site (Figley et al., 2021; Jiang et al., 2020). In contrast, nicotinamide mononucleotide (NMN), a precusor to NAD, can bind to same allosteric pocket to cause a conformational change that relieves autoinhibition and activates SARM1 (Bratkowski et al., 2020; Figley et al., 2021; Zhao et al., 2019). In the long axon projections of neurons, SARM1 activity is thus maintained at a basal level by the activity of the enzyme NMNAT2, which converts NMN to NAD (Coleman & Hoke, 2020). An increase in SARM1 activity, or a loss of NMNAT2 activity, therefore triggers PAxD.

Protein coding variants in *NMNAT2* and *SARM1* have been reported in rare human cases of axonal disorders. Biallelic loss-of-function variants in NMNAT2 have been described in stillborn siblings with Foetal Akinesia Deformation Sequence (Lukacs et al., 2019) and in children with early-onset polyneuropathy (Dingwall et al., 2022; Huppke et al., 2019), with corresponding phenotypes in mouse models providing strong evidence for causality (Gilley et al., 2013; Gilley et al., 2019). In contrast, SARM1 gain-of-function variants have been associated with amyotrophic lateral sclerosis (ALS) (Bloom et al., 2022; Gilley et al., 2021).

Most functionally characterised coding variants in *NMNAT2* are rare and some are associated with highly penetrant axonal diseases. However, the rarity of these variants limits their utility for case–control association analyses in more common disorders. To explore whether more common variation in *NMNAT2* and *SARM1* genes might contribute to susceptibility or act as modifiers in axonal disorders, we examined the effect of promoter variants that could alter *NMNAT2* or *SARM1* expression levels. Identifying regulatory variants with measurable functional effects may expand the spectrum of alleles relevant to risk in the general population. Based on ours and others’ previous findings in mouse neurons lower *NMNAT2* expression could contribute to an individual’s susceptibility to neurodegenerative disease, especially combined with environmental factors or other genetic factors (Ali et al., 2017; Antoniou et al., 2024; Gilley et al., 2019; Loreto et al., 2020; Tribble et al., 2024). For example, individuals exposed to neurotoxic chemotherapeutic agents such as vincristine, known to cause peripheral neuropathy in a subset of patients (Legha, 1986), might be at heightened risk of developing treatment-induced neuropathy if baseline *NMNAT2* expression is reduced. Indeed, there exists broad variation in *NMNAT2* transcript levels in humans, both in post-mortem brain tissue (Ali et al., 2016) and in the retina (Tribble et al., 2024). In parallel, lowering SARM1 levels with antisense oligonucleotides has been shown to protect against axon degeneration in both mouse neurons (Gould et al., 2021) and human neurons (Loreto et al., 2025).

Here, we use functional assays to establish the critical promoter regions required for human *NMNAT2* and *SARM1* expression. We build on published literature exploring the mouse *Nmnat2* promoter (Ljungberg et al., 2012) to show that cAMP signalling also plays a role in human *NMNAT2* expression. We show that naturally occurring single nucleotide changes in the human *NMNAT2* promoter, including one identified in an ALS patient, can reduce promoter activity. Furthermore, we demonstrate that one *SARM1* variant, of relatively common prevalence but evenly distributed between controls and ALS patients, causes a modest increase in promoter activity. These types of promoter region variants could therefore contribute to variable susceptibility of individuals to neurodegenerative disease and may form part of a growing picture of understanding how environmental and genetic factors coalesce on the PAxD pathway, as well as how expression levels of PAxD genes could be modified for therapeutic benefit.

## Materials and Methods

### Cell culture

SH-SY5Y cells (undifferentiated, gifted) and human embryonic kidney (HEK) 293T cells (clone 17, [HEK293T/17]) obtained from the American Type Culture Collection (ATCC, CRL-11268, RRID:CVCL_1926)) were maintained in DMEM with 4,500 mg/L glucose and 110 mg/L sodium pyruvate (Gibco), supplemented with antibiotic-antimycotic solution (penicillin-streptomycin-amphotericin B) and 10% fetal bovine serum. Cells were passaged using TrypLE express enzyme (Thermo Scientific) and not maintained beyond passage 20 for SH-SY5Y cells or passage 30 for HEK293T cells. Cell cultures were incubated at 37°C and 5% CO2. Cell lines were regularly checked for mycoplasma contamination.

### Dibutyryl cAMP treatment

SH-SY5Y cells were treated with 4 mM dibutyryl cAMP (dbcAMP, Sigma-Aldrich / Biosynth) or PBS as a vehicle control, for 24 h.

### Generation of experimental luciferase constructs

The *NMNAT2* and *SARM1* regions 5’ upstream of the translation start site (+1, the ATG start codon) were PCR amplified from SH-SY5Y genomic DNA. The *NMNAT2* and *SARM1* promoter sequences used in this study can be found in the supplementary materials. In *NMNAT2*, 1 variation from the reference genome (GRCh38/hg38 assembly) is present. In *SARM1*, 2 variations from the reference genome (GRCh38/hg38 assembly) are present. These variations are shown in red in the supplementary materials. These are likely due to inherent individual differences present in the genome of the original donor of the cells from which SH-SY5Y cells are derived, or from variations acquired through growing and passaging cells in culture. For *NMNAT2*, BglII and NcoI restriction sites were added to the 5’ and 3’ ends of the promoter regions (-2528, -1929, -1555, -1116, -480, -286, -170 bp relative to the ATG start codon), respectively, using primers and PCR amplification. This allowed for restriction digest-driven insertion of the promoter regions of interest into the pGL4.10 plasmid (Promega), driving the expression of firefly luciferase. In *SARM1* there is an endogenous NcoI restriction site at the translation start site (+1, the ATG start codon). Restriction sites for HindIII were added to the 5’ end of the promoter regions of interest (-1823, -1093, -531, -286 bp), using primers and PCR amplification. As with the *NMNAT2* promoter regions, the *SARM1* promoters were also inserted into the pGL4.10 plasmid using restriction digest and sticky end ligation. Due to the presence of an endogenous NcoI restriction site -1832 bp of the *SARM1* start codon, the -2501 bp *SARM1* promoter plasmid was instead generated by amplifying the region -2501 bp to -667 bp of the *SARM1* start codon, adding a HindIII restriction site at the 5’ end, and taking advantage of an endogenous AflII restriction site (-745 bp from the *SARM1* start codon) to insert the region of interest into an existing *SARM1* promoter pGL4.10 plasmid. Ligated plasmids were amplified in Subcloning Efficiency DH5α Competent Cells (Invitrogen) and plasmid DNA subsequently extracted using the QIAgen Miniprep kit or Midiprep *Plus* kit, according to manufacturer’s instructions.

Site-directed mutagenesis was used to generate single or multiple nucleotide variants in the *NMNAT2* and *SARM1* promoter regions using *PfuUltra* High-Fidelity DNA Polymerase (Agilent Technologies). DpnI restriction digest was used to digest parental (unmutated) DNA and mutated plasmids amplified in MAX Efficiency DH5α Competent Cells (Invitrogen) and extracted as above. Whole plasmid sequencing was performed by Plasmidsaurus using Oxford Nanopore Technology with custom analysis and annotation.

### Luciferase assays

SH-SY5Y or HEK293T cells were seeded onto white, opaque 96-well plates. SH-SY5Y cells were plated at a density of 20 000 cells per well for most luciferase experiments. SH-SY5Y cells were plated at a density of 15 000 cells per well when testing the effects of dbcAMP treatments and when testing the effect of Project MinE variants in the *SARM1* promoter. HEK293T cells were seeded at a density of 15 000 cells per well. The following day, luciferase constructs were introduced into the cells using Lipofectamine 2000 Transfection Reagent (Invitrogen), largely according to the manufacturer’s instructions, although the total volume of DNA-lipid complexes added to each well was 50 µL. 0.02 pmol of pGL4.10 construct (the experimental plasmid driving firefly luciferase expression) and 0.002 pmol of pNL1.1.TK construct (a control plasmid containing the TK promoter driving NanoLuc luciferase) were added to each well (Promega). Prior to performing the luciferase assay, cell medium was changed to a phenol red-free alternative: DMEM with 4,500 mg/L glucose, no glutamine, no phenol red, supplemented with 110 mg/L sodium pyruvate (Gibco), 10% fetal bovine serum and 4 mM L-glutamine. The Nano-Glo Dual-Luciferase Reporter Assay System and a GloMax Explorer plate reader (Promega) was used to measure luminescent signal 24 h after transfection, according to the manufacturer’s instructions. In the dbcAMP experiments, drug treatment was administered 24 h after transfection and luminescent signal measured 24 h following that. Background-subtracted firefly luciferase activity is reported relative to the activity of background-subtracted NanoLuc luciferase, which is then normalised to the luminescence readout from cells transfected with promoterless pGL4.10, for each biological replicate. Three technical replicates (wells) were averaged for each independent experiment.

### Quantitative reverse transcriptase PCR (qRT-PCR)

700 000 SH-SY5Y cells per well were seeded in 6-well plates. After 24 h they were treated with dbcAMP or PBS as a vehicle control. 24 h after treatment, cells were washed once with PBS, detached using TrypLE express enzyme (Thermo Scientific) and centrifuged to pellet (500 x *g* for 5 minutes). The pellet was washed once in PBS and centrifuged at 300 x *g* for 5 minutes to re-pellet. RNA was extracted from the cells using the QIAgen RNeasy kit (mini), according to the manufacturer’s instructions and including the optional on-column DNase I digestion. 1 µg of RNA was reverse transcribed into cDNA using the QIAgen QuantiTect Reverse Transcription kit, according to the manufacturer’s instructions and using a T100 thermal cycler (Bio-Rad). No reverse transcriptase (No RT) controls were included for each sample. 1 µL of the 20 µL reverse transcription reaction mix was directly used for each quantitative PCR reaction. For each experimental condition the average was taken from three technical replicates. Quantitative PCR was performed in white hard-shell 96-well PCR plates (Bio-Rad HSP9655) using the CFX96 Touch Real-Time PCR Detection System, iTaq Universal SYBR Green Supermix (Bio-Rad) and 500 nM final concentration of each of the primers. The primers were as follows: *NMNAT2:* forward 5’-GATTGGATCAGGGTGGACC -3’, reverse 5’- TCCGATCACAGGTGTCATGG -3’ (150 bp product) (Buonvicino et al., 2018), *ACTB: forward 5’:* ACAGAGCCTCGCCTTTGC -3’, reverse 5’- CGCGGCGATATCATCATCCA -3’ (76 bp product), *IPO8*: forward 5’-GGCATACAGTTTAACCTGCCAC -3’, reverse 5’- CAGGAGAGGCAT CATGTCTGTAA -3’ (118 bp product) (Rácz et al., 2021). The PCR conditions were 95°C for 30 seconds then 45 x (95°C for 5 seconds, 60°C for 15 seconds). RNA extraction, reverse transcription and quantitative PCR were all performed in the same day to avoid freeze-thawing of the samples. Primer efficiencies were determined from a serial dilution of cDNA pooled from all the sample groups of one replicate and calculated to be 103% (*NMNAT2*), 100% (*ACTB*) and 91% (*IPO8*) (Supp. Fig. 1A-C). To confirm amplification of a single target for each primer pair, melt curve analysis was performed by increasing 0.5°C every 5 seconds from 65°C to 95°C. qRT-PCR products were run on a 1% agarose gel alongside a 1 Kb Plus DNA ladder (Invitrogen) to further assess primer specificity (Supp. Fig. 1D). Data were analysed using the 2^ΔΔCt^ method relative to the housekeeping genes *ACTB* and *IPO8* (data shown for both) and the ΔCt of vehicle treated cells. Within each replicate, fold change was normalised to *NMNAT2* expression in vehicle treated cells (so that *NMNAT2* expression in vehicle treated cells always equals 1).

### Sequence alignment

The translation start codon and 286 base pairs 5’ of *NMNAT2* for mouse (*mus musculus*), rat (*rattus norvegicus*) and human (*homo sapiens*) were aligned using Clustal Omega (1.2.4) Multiple Sequence Alignment (Madeira et al., 2024).

### *In silico* analysis of candidate regulatory elements within the human *NMNAT2* promoter

To identify potential regulatory elements upstream of the characterized CRE2 element, a 116 bp interval, −286 to −170 bp relative to the *NMNAT2* translation start site (Chr1:183,418,553–183,418,438, GRCh38/hg38 assembly, minus strand) was retrieved and screened for repetitive and low-complexity sequence using RepeatMasker (v4.1.2; Dfam v3.7) (http://www.repeatmasker.org/). *De novo* motif discovery was then performed using MEME Suite v5.5.4 (Bailey et al., 2015) (motif width 6–12 bp; ZOOPS model; human genomic background). Motif occurrences and statistical support for positional clustering were evaluated using MAST (Bailey & Gribskov, 1998).

To infer potential transcription factor binding activities, discovered motifs were compared against curated transcription factor binding profiles in JASPAR CORE and TRANSFAC-like databases using STAMP (Mahony & Benos, 2007) and matches with E-values < 1 × 10⁻² were considered for functional annotation.

### Variant screening in population databases

#### GnomAD

To identify naturally occurring variants within the functional CRE2 element, we queried the Genome Aggregation Database (gnomAD) version 4.1.0 (Karczewski et al., 2020; https://gnomad.broadinstitute.org/) for all variants within Chr1:183,418,403–183,418,414 (GRCh38/hg38), corresponding to the 12 bp CRE2 sequence in the *NMNAT2* promoter. Variant allele counts were extracted from the gnomAD browser interface.

#### Project MinE

We obtained variant-level data from Project MinE data freeze 1 (DF1), comprising 4,366 individuals with ALS and 1,832 age- and sex-matched controls (van der Spek et al., 2019). For *NMNAT2*, we interrogated all variants located within Chr1:183,418,268–183,418,602 (GRCh38/hg38), corresponding to the 335 bp upstream of the translation start site and annotated by Project MinE as “5′ UTR variants”. No variants within this interval were annotated as ‘transcription factor binding site variants’ or ‘upstream gene variants’. Note that in the Project MinE database DF1 the *NMNAT2* gene is annotated only as far as 335 bp 5’ of the ATG start codon. For *SARM1*, we extracted all variants annotated as ‘5′ UTR variants’, ‘transcription factor binding site variants’, or ‘upstream gene variants’ within our defined promoter region (Chr17:28,369,532–28,372,032; 2501 bp upstream of the translation start site). To prioritise *SARM1* candidates for functional testing, we selected variants showing >1.4-fold enrichment in either cases or controls, as well as variants for which three or more alleles were observed exclusively in one group. We also selected the four most common *SARM1* variants, irrespective of whether they showed enrichment in case or control groups. All *NMNAT2* and the prioritised *SARM1* variants were subsequently subjected to experimental evaluation.

#### Statistics

Data are presented as mean ± SEM. Statistical analyses were performed using GraphPad Prism 10.2.1. Statistical significance was set at *p* ≤ 0.05. The Shapiro-Wilk test was used to assess whether data sets are normally distributed and parametric or non-parametric statistical analyses applied accordingly. The statistical tests used and N numbers for each data set are reported in the figure legends.

## Results

### 286 base pairs drive human *NMNAT2* promoter activity

We first set out to determine the region 5’ of the translation start site (ATG codon, +1) of human *NMNAT2* required for maximal gene expression. To this end we cloned different portions of the relevant DNA and used luciferase assays to measure their effects on transcription. We used two, easy to transfect, cells lines: SH-SY5Y (a cell line of human neuroblastoma origin) and HEK293T cells (of human embryonic kidney origin but with some neuron-like properties (He & Soderlund, 2010; Shaw et al., 2002)). In both cell lines, maximal *NMNAT2* promoter activity is driven by the first 286 bp upstream of the ATG start codon (Fig. 1).

**Figure 1:**
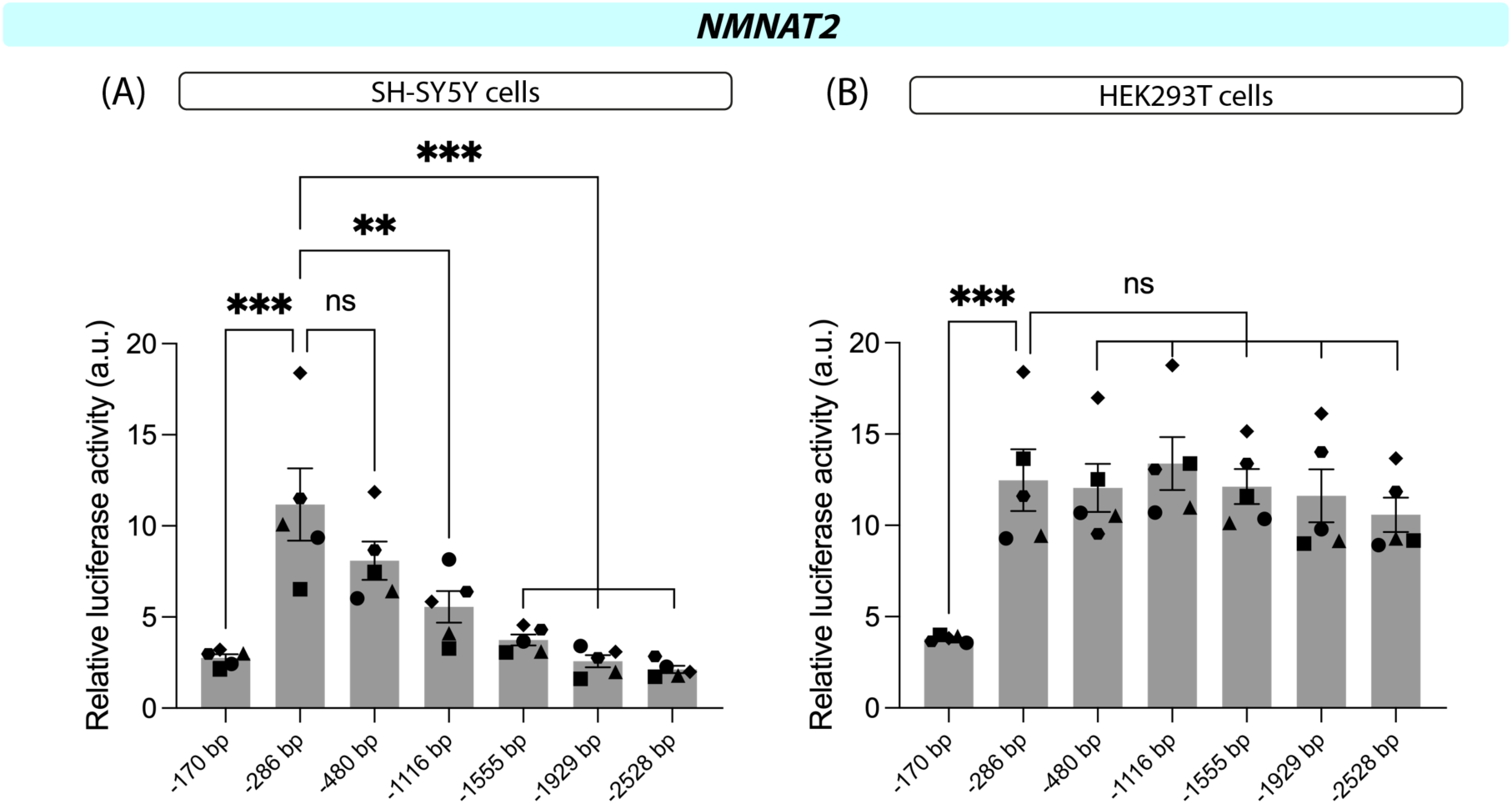
286 base pairs drive maximal expression of human *NMNAT2*. Luciferase assays in SH-SY5Y cells (A) and HEK293T cells (B) comparing various lengths of the *NMNAT2* promoter region suggest that the first 286 base pairs (bp) upstream (5’) of the canonical translation start site (ATG) drive human *NMNAT2* expression. However, the region -286 to -2528 appears to increasingly repress *NMNAT2* expression, specifically in SH-SY5Y cells (A). Data are presented as mean ± SEM. N = 5 (independent experiments are indicated by different shape data points). Statistical significance was determined by one-way ANOVA: (A) *F* (6,28) = 13.25, *P* < 0.001. (B) *F* (6,28) = 6.891, *P* < 0.001. The statistical significance indicated on the graphs (* *P* ≤ 0.05, ** *P* ≤ 0.01, *** *P* ≤ 0.001) was determined by Dunnett’s multiple comparisons tests. For clarity, only statistical significance compared to -286 bp is shown.

Intriguingly, as the sequence is extended in the 5’ direction, we see an accumulative repression of promoter activity specifically in SH-SY5Y cells, so that when 2528 bp are upstream of our luciferase reporter, relative activity (a reflection of luciferase expression and therefore promoter activity) is comparable to the activity when luciferase expression is driven by 170 bp of the region upstream of the *NMNAT2* ATG start codon (Fig. 1A). In contrast, in HEK293T cells, relative activity of the luciferase reporter remains consistent regardless of how far the promoter region is extended beyond 286 bp (Fig. 1B).

### The human *NMNAT2* promoter contains one cAMP response element

A study on the mouse *NMNAT2* promoter identified two cAMP response elements (CREs) (Ljungberg et al., 2012). CREs are consensus sequences that allow the binding of the transcription factor CRE-binding protein (CREB) and its family members in order to regulate gene expression (Mayr & Montminy, 2001; Sakamoto et al., 2011). We aligned the 286 bp 5’ of the ATG start codon in human, mouse and rat allowing us to predict where the two CREs found in mice would lie if conserved in the human sequence (Fig. 2). CRE2, the site most proximal to the translation start site, was completely conserved, bar one flanking base, across the three species. However, CRE1, while identical between mouse and rat, was poorly conserved to humans (Fig. 2). In particular, the core transcription factor binding residues TGACG are not well conserved in CRE1 suggesting this site is unlikely to be functional in human.

**Figure 2:**
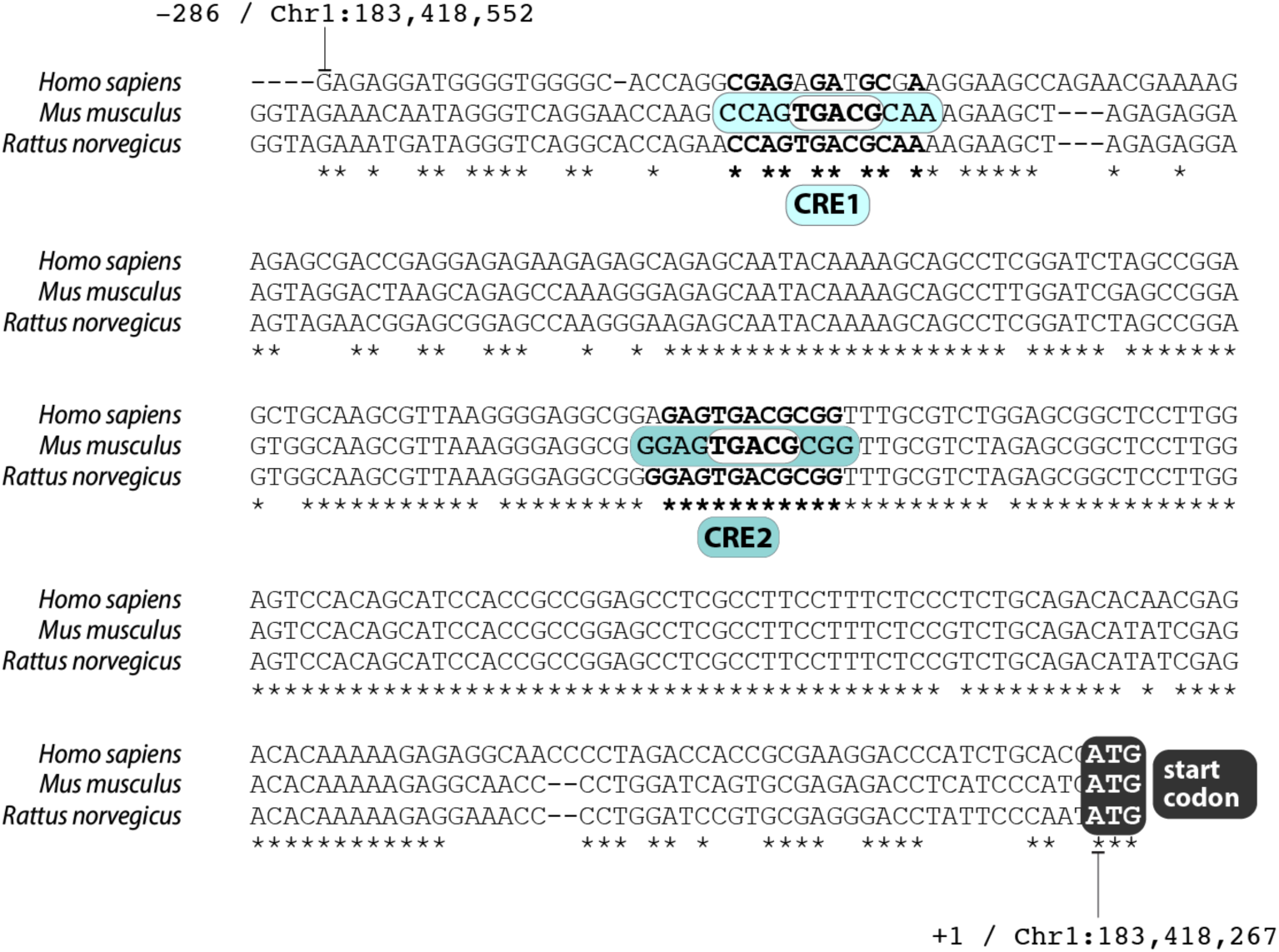
One cAMP response element (CRE) in the *NMNAT2* promoter is conserved from mouse to human. Sequence alignment of the 286 base pairs (bp) 5’ of the translation start site (ATG) of *NMNAT2* in mouse (*Mus musculus*), rat (*Rattus norvegicus*) and human (*Homo sapiens*) shows that cAMP response element 2 (CRE2) (Ljungberg et al., 2012) is well conserved whereas CRE1 is less so. TGACG is the core CRE binding half-site. Asterisks indicate conserved bases. Chromosomal coordinates for the human sequence are indicated (GRCh38/hg38).

To test whether these prospective CREs were functional in the human gene we mutated four bp in each of the regions (Fig. 3A). Unsurprisingly, mutating CRE1, the poorly conserved site, has no effect on promoter activity in SH-SY5Y cells (Fig. 3B). However, mutating CRE2, the evolutionarily conserved site, causes a substantial reduction in promoter activity, so that the luciferase activity drops from 17.4 ± 2.1 in control (-286 bp) to 5.0 ± 0.2 (*P* < 0.001. Fig. 3B). Mutating both CRE1 and CRE2 in tandem (i.e. within the same construct) produces the same effect as mutating CRE2 alone, indeed providing further evidence that the poorly conserved CRE1 sequence in the human *NMNAT2* promoter has no major influence on gene expression (Fig. 3B).

**Figure 3:**
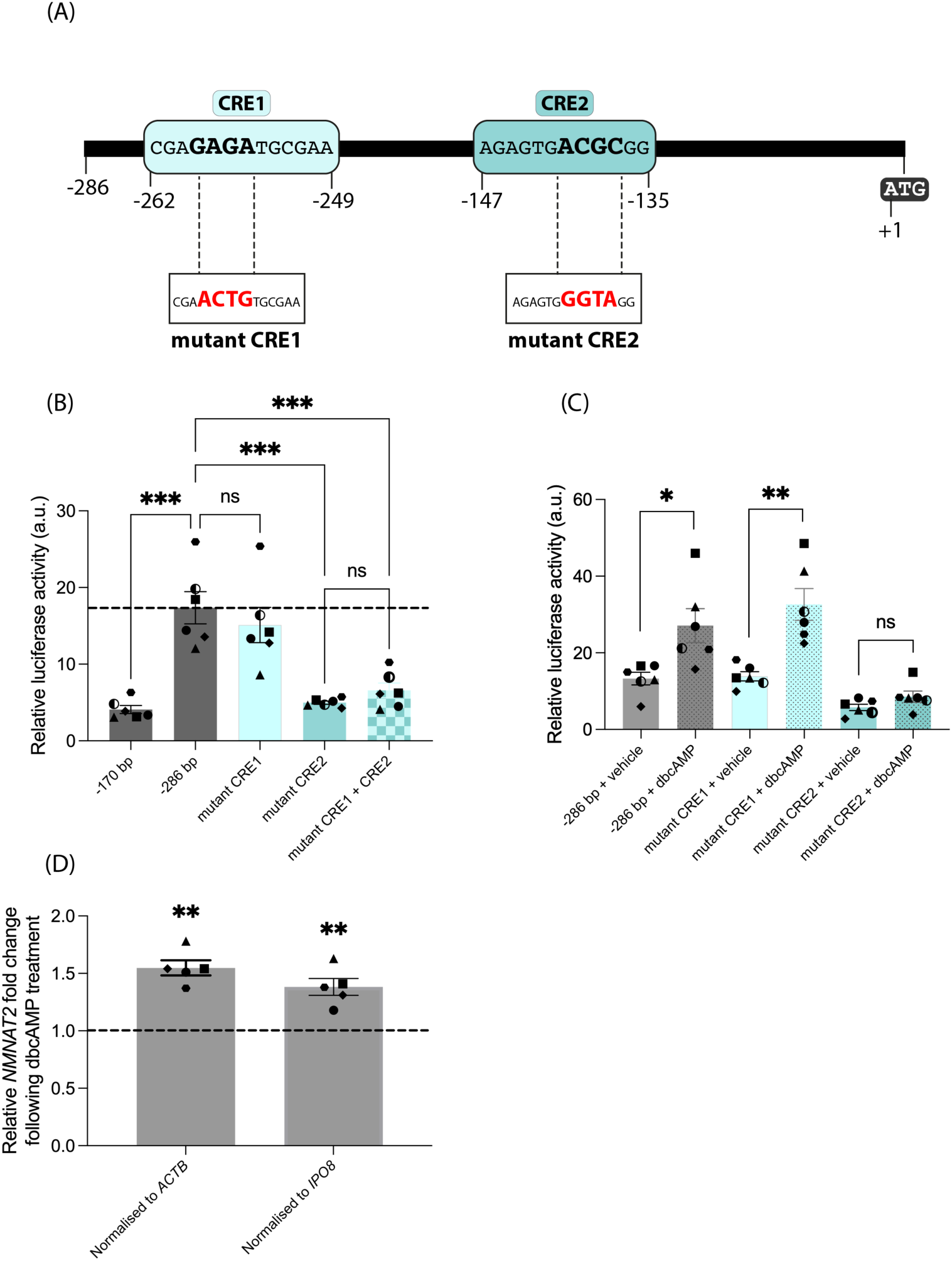
Human *NMNAT2* promoter activity is upregulated by cAMP, acting through cAMP response element 2 (CRE2). (A) Schematic showing mutations made to predicted cAMP response elements 1 and 2 (CRE1 and CRE2). (B) Luciferase assays in SH-SY5Y cells demonstrate that mutating CRE1 has no effect on human *NMNAT2* promoter activity, whereas mutation in CRE2 reduces promoter activity. Statistical significance was determined by one-way ANOVA: *F* (4,25) = 17.35, *P* < 0.001. Selected comparisons, determined by Tukey’s multiple comparisons tests, are shown on the graph. (C) Luciferase assays in SH-SY5Y cells show that 24 h treatment with 4 mM dbcAMP increases human *NMNAT2* promoter activity. When CRE2 is mutated the effect of dbcAMP is abolished. Statistical significance was determined by unpaired t test, corrected for multiple comparisons using the Holm-Šidák method. (D) qRT-PCRs show that 24 h treatment with 4 mM dbcAMP increases endogenous *NMNAT2* expression in SH-SY5Y cells. *NMNAT2* expression is normalised to the housekeeping genes *ACTB* and *IPO8*. The statistical significance was determined by one sample t tests comparing to a value of 1. Data are presented as mean ± SEM. N = 5-6 (independent experiments are indicated by different shape data points). * *P* ≤ 0.05, ** *P* ≤ 0.01, *** *P* ≤ 0.001.

To test whether the *NMNAT2* promoter indeed responds to changes in cAMP levels, we treated SH-SY5Y cells expressing luciferase under the control of the *NMNAT2* promoters with 4 mM dbcAMP for 24 h, or PBS as a vehicle control. When cells expressing luciferase driven by wildtype or mutant CRE1 *NMNAT2* promoters are treated with dbcAMP an increase in luciferase activity is measured, from 13.3 ± 1.6 to 27.1 ± 4.4 (wildtype, *P* = 0.03) and from 13.9 ± 1.2 to 32.6 ± 4.2 (mutant CRE1, *P* = 0.004) (Fig. 3C). However, when CRE2 is mutated the effect of dbcAMP is abolished, so that when treated with vehicle the relative luciferase activity measured is 5.8 ± 0.8 and when treated with dbcAMP activity is 8.6 ± 1.5 (*P* = 0.13) (Fig. 3C).

We next sought to test whether treatment with dbcAMP increases endogenous *NMNAT2* expression in SH-SY5Y cells. Quality controls confirmed primer efficiency (Supp. Fig. 1A-C) and the generation of one PCR product per primer pair (Supp. Fig. 1D). We normalised *NMNAT2* expression to two different housekeeping genes and found that treatment with dbcAMP for 24 h increases SH-SY5Y cell *NMNAT2* Mrna 1.5 ± 0.07 fold when normalised to *ACTB* (*P* = 0.0011) and 1.4 ± 0.07 fold when normalised to *IPO8* (*P* = 0.0066) (Fig. 3D). This confirms that human *NMNAT2* expression responds to changes in cAMP levels.

### Further regulatory features exist within the *NMNAT2* promoter

While CRE2 appears to be a major driver of human *NMNAT2* promoter activity, our data suggests that it is not the only key element driving *NMNAT2* expression. CRE2 is found within the first 170 bp upstream of the *NMNAT2* translation start codon (Fig. 3A), and while mutating this site drastically reduces promoter activity (Fig. 3B), expression of CRE2 alone (in the case of the -170 bp construct), is not sufficient for maximal promoter activity (Fig. 1 and Fig. 3B). It is thus likely that CRE binding transcription factors may interact with other transcription factors binding in the region between 170 and 286 bp upstream of the start codon to regulate *NMNAT2* expression. We therefore undertook computational and functional approaches to characterize potential regulatory elements within this 116 bp interval.

RepeatMasker analysis revealed that 47% (54 bp) of the −286 to −170 bp interval upstream of the *NMNAT2* translation start site consists of AT-rich low-complexity sequence. These regions were masked prior to motif discovery to reduce compositional bias.

*De novo* motif analysis identified three short motifs (8–9 bp; MEME1–3), none of which demonstrated strong statistical enrichment (Table 1 and Supplementary Fig. 2A). Although MAST detected multiple instances of these motifs within the interval (combined motif-set *P* = 1.4 × 10⁻⁵), motif occurrences were spatially dispersed and lacked positional clustering or organization consistent with cooperative cis-regulatory modules. The modest MAST P value likely reflects the short, degenerate nature of the motifs rather than biologically meaningful regulatory specificity.

**Table 1:** ‘MEME’ motifs in the human *NMNAT2* promoter. IUPAC ambiguity codes: R = A/G; Y = C/T; K = G/T; S = G/C; W = A/T; M = A/C; N = any base. TF = transcription factor.

| **Motif** | **Consensus sequence** | **Predicted human / mammalian TF families (STAMP similarity)** | **Representative human TFs in family** | **Notes** |
| --- | --- | --- | --- | --- |
| **MEME1** | ATASARARG | KLF family (C2H2 zinc-finger); HMG-box SOX family | KLF4, SOX9, HMGIY | SOX9 physically interacts with CREB in osteochondrogenesis (Zhao et al., 2009). |
| **MEME2** | YCGGAKCT | bHLH family; ETS transcription factors; C2H2 zinc-finger | NHLH1, ELK4, ZNF354C | bHLH acts as a regulator in neuronal differentiation, reprogramming, and cell fate determination (Nelms & Labosky, 2010).<br>ETS member proteins regulate genes associated with neuronal differentiation, nervous system development, axon, and synaptic regulation (Babal et al., 2023). |
| **MEME3** | GSSTGGKG | CREB/ATF family; REST/NRSF; TFAP2A; NFIC | CREB1, REST, TFAP2A, NFIC | CREB1 interaction is relevant as CRE2 is a CRE motif. REST binding is supported by ENCODE ChIP-seq data. |

Comparison of the three discovered motifs to curated transcription factor binding profiles using STAMP yielded weak similarities to motifs associated with KLF4, HMG-IY and SOX9 (MEME1); basic helix–loop–helix (bHLH) and ETS family members, including NHLH1 and ELK4 (MEME2); and REST, TFAP2A and CREB1-like factors (MEME3) (Table 1).

We subsequently mutated the possible transcription factor binding sites (and one site detected slightly further upstream, MEME3-1) and tested the effect of mutation on promoter activity in SH-SY5Y cells (Supp. Fig. 2A-B). For consistency, all the mutated motifs were introduced into the -483 bp *NMNAT2* promoter, since sites MEME1-1 and MEME3-1 are found further upstream than -286 bp (Supp. Fig. 2A), precluding introduction into the -286 bp promoter construct. None of the mutations of these candidate sites elicited the hypothesized reduction in promoter activity. In fact, mutation at one site (MEME2-2) appears to produce a modest increase in promoter activity, suggesting this motif may be involved with repressive transcription factor activity. Nevertheless, overall, our systematic computational and experimental analysis provides limited evidence for discrete transcription factor binding sites within the -286 to -170 bp region that are individually essential for *NMNAT2* promoter activity.

### Naturally occurring variants in the cAMP response element reduce *NMNAT2* promoter activity

Having identified CRE2 as a critical regulatory element driving human *NMNAT2* expression, we queried gnomAD v4.1.0 (Karczewski et al., 2020) to determine whether naturally occurring variants are present within this region (Chr1:183,418,403–183,418,414, GRCh38/hg38). Eight variants (GNO1-GNO8) were identified within CRE2, all of which exhibit very low allele frequencies (Fig. 4A). We individually mutated each of these sites to reflect the naturally occurring human variants (Fig. 4B) and tested their effect on *NMNAT2* promoter activity in SH-SY5Y cells. Mutating a single nucleotide is sufficient to reduce promoter activity for four out of the eight naturally occurring human variants, GNO4, GNO5, GNO6 and GNO7 (Fig. 4C). Notably, three of these variants (GNO4, GNO5 and GNO6) lie directly within the predicted transcription factor binding region (the core half-site TGACG, bold in Fig. 4B) (Sakamoto et al., 2011). These results demonstrate that natural variation in *NMNAT2* promoter activity does exist within the human population, albeit rare. Individuals carrying these single nucleotide variants (SNVs) may exhibit reduced *NMNAT2* expression, potentially increasing susceptibility to PAxD-mediated axon degeneration. We thus next hypothesised that SNVs lowering expression from the *NMNAT2* promoter may be enriched in neurodegenerative disease populations.

**Figure 4:**
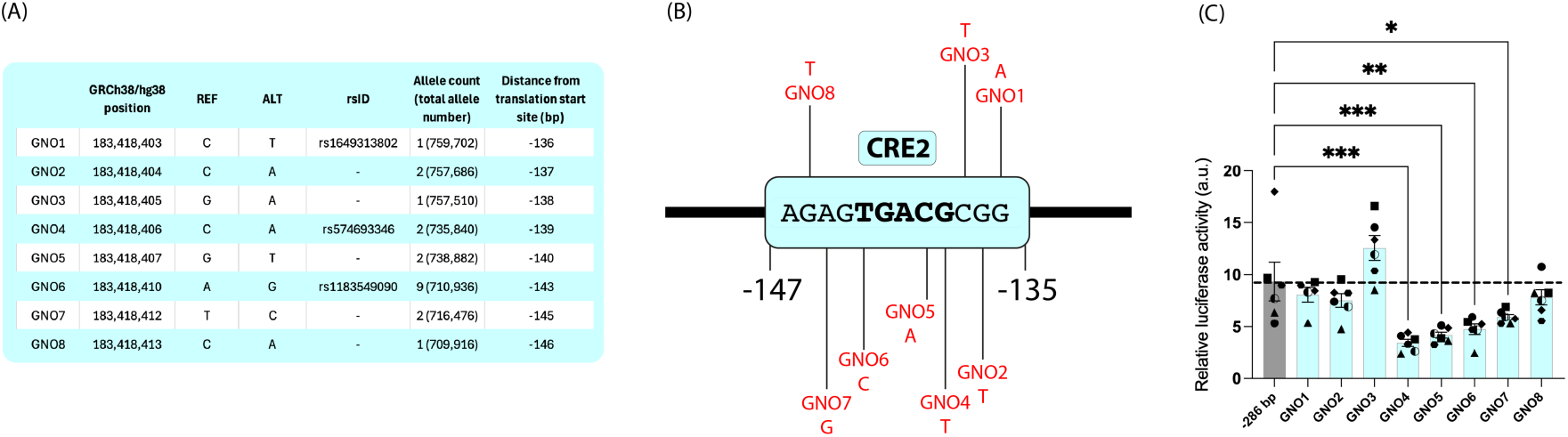
Naturally occurring single nucleotide variants (SNVs) in cAMP response element 2 (CRE2) reduce *NMNAT2* promoter activity. (A) Eight naturally occurring SNVs are found in the CRE2 region in the gnomAD database. REF = reference allele, ALT = alternate allele, bp = base pairs. (B) Schematic showing the position of the eight gnomAD SNVs in the CRE2 region. Note: In humans *NMNAT2* is found on the minus strand. Reference (REF) and alternate (ALT) bases in (A) are relative to the plus strand, whereas the CRE2 sequence shown in (B) is the minus strand promoter sequence of the *NMNAT2* promoter. The bold sequence represents the predicted transcription factor binding half-site. (C) Luciferase assays in SH-SY5Y cells show that four of the eight SNVs in CRE2 cause a reduction in human *NMNAT2* promoter activity. Data are presented as mean ± SEM. N = 6 (independent experiments are indicated by different shape data points). Statistical significance was determined by one-way ANOVA: *F* (8,44) = 10.83, *P* < 0.001. The statistical significance indicated on the graph (* *P* ≤ 0.05, ** *P* ≤ 0.01, *** *P* ≤ 0.001) was determined by Dunnett’s multiple comparisons tests.

### A single nucleotide variant (SNV) in an ALS patient reduces *NMNAT2* promoter activity

To this end, we used the Project MinE database data freeze 1 (DF1) (van der Spek et al., 2019), which contains 4,366 sporadic amyotrophic lateral sclerosis (ALS) cases and 1,832 age and sex matched controls. Within this dataset, 12 SNVs are annotated within the *NMNAT2* promoter region (Chr1:183,418,268–183,418,602; 335 bp upstream of the ATG) (Fig. 5A). None of the functional CRE2 element variants found in gnomAD (Fig. 4) are found in Project MinE DF1. Of the 12 Project MinE SNVs, 10 are found exclusively in the ALS population, 1 only in controls and 1 in both ALS cases and controls, however all variants, as with those found in CRE2 in gnomAD, are extremely rare (Fig. 4A and 5A). We subsequently introduced the 12 SNVs from Project MinE (PMV1-12) into the human *NMNAT2* promoter. Depending on the location of the variant, the appropriate control comparison was either 286 bp upstream of the start codon (Fig. 5B) or 480 bp upstream of the start codon (Fig. 5C). Our luciferase reporter assay suggests that one Project MinE variant (PMV6) reduces promoter activity in SH-SY5Y cells (Fig. 5B). Luciferase activity dropped from 11.0 ± 1.7 in controls to 4.9 ± 0.5 with PMV6 (*P* = 0.04). This presents the possibility that the individual in the Project MinE database who carries PMV6 could have lower expression of *NMNAT2* at the transcriptional level which could contribute to disease pathogenesis by facilitating SARM1 activation.

**Figure 5:**
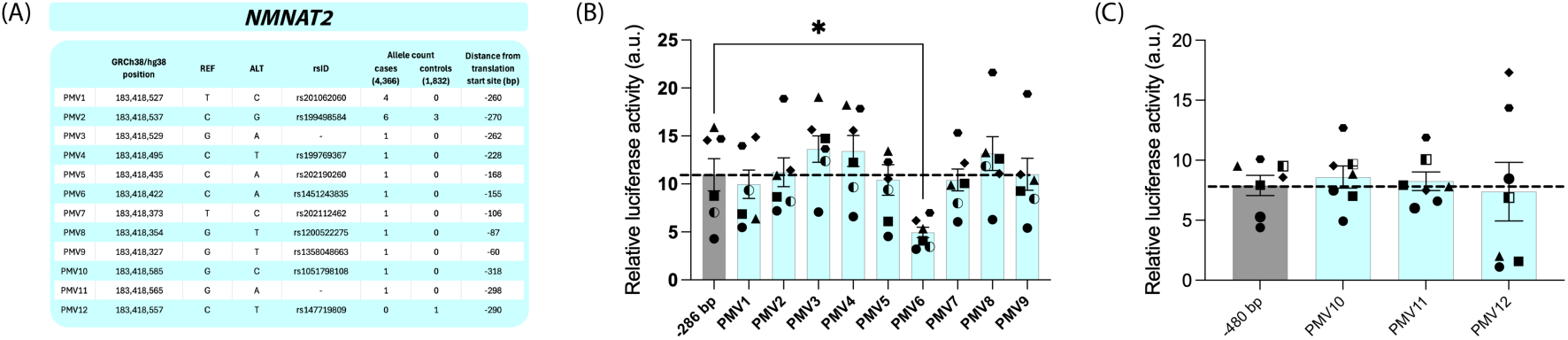
A single nucleotide variant (SNV) found in an ALS patient reduces *NMNAT2* promoter activity. (A) 12 SNVs are found in the *NMNAT2* promoter region in the Project MinE database (data freeze 1, DF1). PMV = Project MinE Variant, REF = reference allele, ALT = alternate allele, bp = base pairs. (B) Luciferase assays in SH-SY5Y cells suggest PMV6 reduces *NMNAT2* promoter activity. (C) Luciferase assays in SH-SY5Y cells suggest PMV10, PMV11 and PMV12 have no effect on *NMNAT2* promoter activity. Data are presented as mean ± SEM. N = 7 (independent experiments are indicated by different shape data points). Statistical significance was determined by one-way ANOVA: (B) *F* (9,60) = 2.859, *P* = 0.007, (C) *F* (3,24) = 0.1312, *P* = 0.94. The statistical significance indicated on the graphs (* *P* ≤ 0.05, ** *P* ≤ 0.01, *** *P* ≤ 0.001) was determined by Dunnett’s multiple comparisons tests.

### 1093 base pairs drive human *SARM1* promoter activity

We next set out to use a similar experimental approach to probe the human *SARM1* promoter. The SARM1 enzyme acts downstream of NMNAT2, and when activated causes axon death. Gain-of-function mutations in the coding region of SARM1 cause it to be hyperactive and have been linked to ALS (Bloom et al., 2022; Gilley et al., 2021). In both cell lines examined, 286 or 531 bp immediately upstream of the *SARM1* start codon are insufficient to robustly drive *SARM1* expression (Fig. 6). In both cell types, maximal promoter activity appears to be elicited with 1093 bp, which is retained as the promoter region is extended to 2501 bp (Fig. 6).

**Figure 6:**
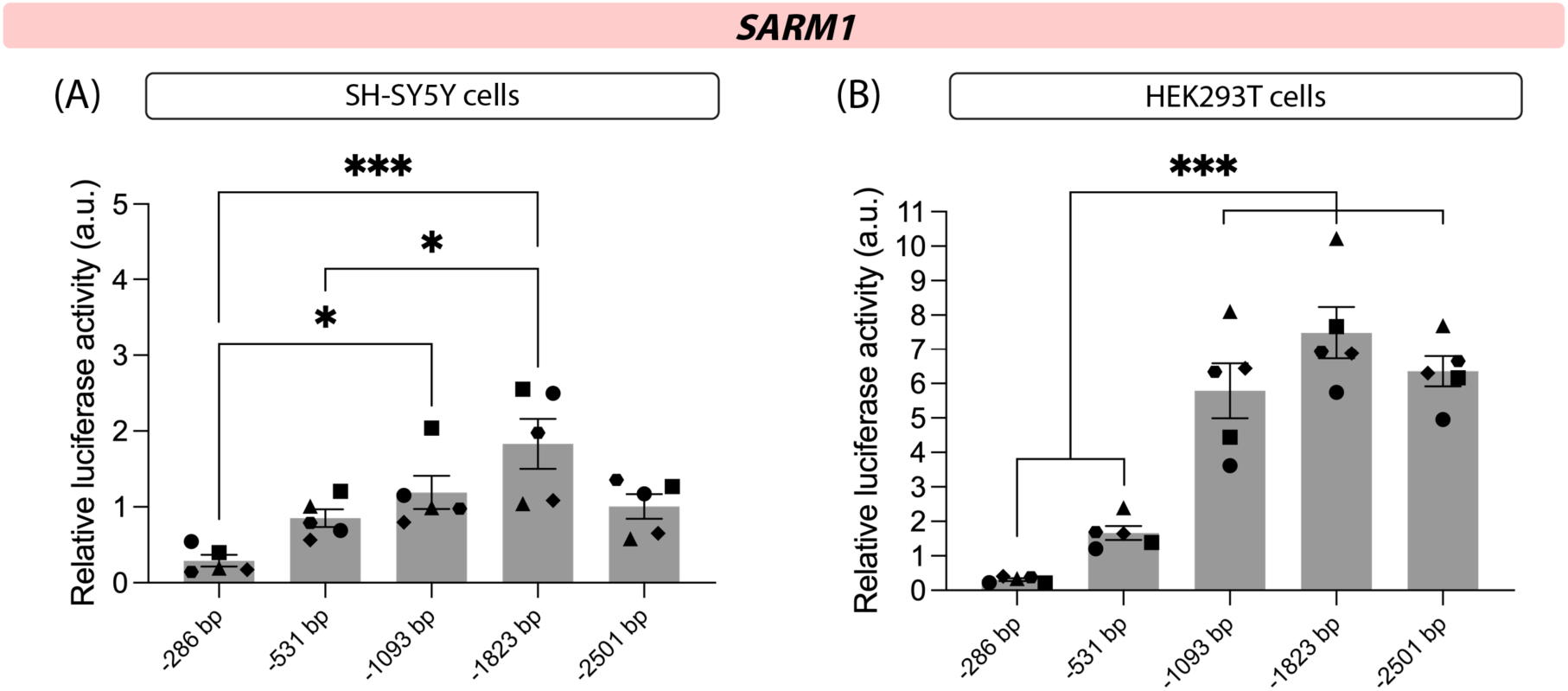
A 1093 base pair promoter drives full expression of human *SARM1*. Luciferase assays in SH-SY5Y cells (A) and HEK293T cells (B) comparing various lengths of the *SARM1* promoter suggest that the first 1093 base pairs (bp) upstream (5’) of the canonical translational start site (ATG) drive human *SARM1* expression. Data are presented as mean ± SEM. N = 5 (independent experiments are indicated by different shape data points). Statistical significance was determined by one-way ANOVA: (A) *F* (4,20) = 7.714, *P* < 0.001, (B) *F* (4,20) = 34.45, *P* < 0.001. The statistical significance indicated on the graphs (* *P* ≤ 0.05, ** *P* ≤ 0.01, *** *P* ≤ 0.001) was determined by Tukey’s multiple comparisons tests.

### A common single nucleotide variant (SNV) found in patients and controls increases *SARM1* promoter activity

As with *NMNAT2*, we probed the Project MinE database to examine whether the *SARM1* promoter region (Chr17:28,369,532-28,372,032, 2501 bp) contains human variants. Since we were able to examine a larger region (2501 bp) upstream of the *SARM1* start codon (compared to only 335 bp for *NMNAT2*) we stratified the variants to examine in our functional assays in order to test variants enriched in either the ALS population or the control population, or to test the most commonly found variants regardless of subgroup enrichment (Fig. 7A, our filtering method is described in more detail in our Materials and Methods). Most of the 16 *SARM1* variants we chose to study have no effect on promoter activity in SH-SY5Y cells (Fig. 7B-C). One variant, PMV13, shows a modest but significant increase in promoter activity from 3.1 ± 0.5 in control to 4.4 ± 0.4 in PMV13 (*P* = 0.005, Fig. 7B). Interestingly, this variant is not enriched in either control or patient population (Fig. 7A) suggesting it does not contribute to risk of ALS. However, with a relatively high number of occurrences in the database we probed, this variant does present a common single nucleotide polymorphism (SNP) with the potential to modify *SARM1* expression, which could be used in future disease-association studies.

**Figure 7:**
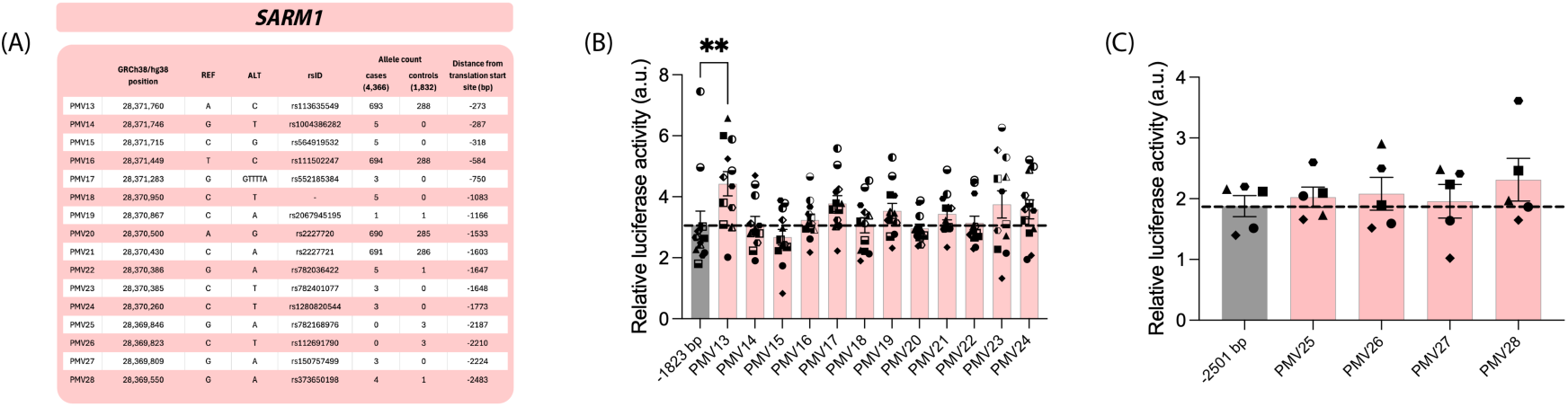
A single nucleotide variant (SNV) increases *SARM1* promoter activity. (A) Many variants are found in the *SARM1* promoter region in the Project MinE database (data freeze 1, DF1), of which 16 were selected for functional testing, as per the criteria outlined in Materials and Methods. PMV = Project MinE Variant, REF = reference allele, ALT = alternate allele, bp = base pairs. (B) Luciferase assays in SH-SY5Y cells suggest that PMV13 increases *SARM1* promoter activity. Statistical significance was determined by Kruskal-Wallis (*P* = 0.009). The statistical significance indicated on the graph (* *P* ≤ 0.05, ** *P* ≤ 0.01, *** *P* ≤ 0.001) was determined by Dunn’s multiple comparisons tests. (C) Luciferase assays in SH-SY5Y cells suggest that PMVs 25-28 have no effect on *SARM1* promoter activity. Data are presented as mean ± SEM. N = 5-12 (independent experiments are indicated by different shape data points). Statistical significance was determined by one-way ANOVA (*F* = (4,20) = 0.4094, *P* = 0.80).

## Discussion

In our study we use functional, in-cell assays to identify critical genomic regions for the human *NMNAT2* and *SARM1* promoters and to explore the effect of naturally occurring variations within these regions, found in the general population and in patients with neurodegenerative disease.

For *NMNAT2*, very minimal promoter activity is recorded from the first 170 bp upstream of the translation start codon (ATG). This is perhaps unsurprising, since according to annotation on the genome browser Ensembl, 170 bp contains just 57 bp in addition to the 5’UTR. Instead, the first 286 bp 5’ from the translation start codon (ATG) appears to robustly drive gene expression. We establish that while the mouse *NMNAT2* promoter requires two cAMP response elements (CREs) (Ljungberg et al., 2012), the human promoter appears to rely on just one, through which human *NMNAT2* levels can be increased by cAMP. Our data also suggests further regulatory elements exist within the -286 to -170 bp interval of the *NMNAT2* promoter, although our computational and experimental approaches suggest that functional elements within this region may act in synergy to elicit an effect on promoter activity, rather than possessing discrete transcription factor binding sites. Naturally occurring SNVs in the CRE exist in the human population and these have the ability to lower *NMNAT2* promoter activity; our assays suggest that in the case of some variants this can be by greater than 60%. Evidence from mice suggests that low *NMNAT2* expression close to this level increases susceptibility to axon degeneration or cell death by, for example, vincristine or the mitochondrial toxin CCCP (Ali et al., 2017; Gilley et al., 2019; Loreto et al., 2020; Tribble et al., 2024). Furthermore, our probing of the ALS database Project MinE identifies one variant in the *NMNAT2* promoter (PMV6) that reduces promoter activity by more than 50%. This variant is found only in an ALS patient and absent from controls in Project MinE DF1. Notably, the promoter variants in *NMNAT2* are extremely rare. PMV6, for example, has an allele count of just 4 in gnomAD v4.1.0 (Karczewski et al., 2020).

A repression of *NMNAT2* promoter activity occurs from -286 bp to -2528 bp, which is cell-type specific. One transcription factor that could mediate such an effect is the repressor element 1 (RE1) silencing transcription factor (REST, also known as neuron-restrictive silencer factor, NRSF, and X2 box repressor, XBR). REST has been shown to be important for development and function of the nervous system (Jin et al., 2023). Our own and others’ (Chang et al., 2025) bioinformatics analysis, as well as ChiPSeq experiments (Johnson et al., 2007), have indicated that REST can interact with the human *NMNAT2* promoter. Chang et al. (2025) also suggest that the transcription factors ATF4 or SOX11 could act as *NMNAT2* transcriptional repressors. Notably, ATF4 belongs to the ATF/CREB family of transcription factors which can bind at CREs (Ameri & Harris, 2008), a regulatory feature we show to be present in the human *NMNAT2* promoter, although in our experimental conditions we find that CRE2 in humans is necessary to drive transcriptional activity, rather than repress it. Probing further into this repressive region could reveal targets for the manipulation of *NMNAT2* transcriptional activity; for example relieving inhibition to increase the expression of *NMNAT2* and promote axonal survival.

For *SARM1* we find that 1093 bp upstream of the translation start codon are required for full promoter activity. Extending the promoter further, up to 2501 bp 5’ of the start codon, has little effect, neither increasing nor decreasing the promoter’s activity. Some rare variants in the promoter region are found exclusively in ALS patients, or exclusively in controls, in Project MinE DF1. More common variants are also found, but these appear to be equally distributed between controls and patients. For example, PMV13 is not enriched in ALS patients or in controls, but it is relatively common with an allele frequency of 0.08 in both groups. In our luciferase assay, we found that PMV13 causes a statistically significant increase in promoter activity. Our experiments transiently express the promoter variants in front of a reporter, so we should be mindful to consider that the effects may differ from the same variant fully integrated into the genome. The transcriptional regulation probed in our assays does not take into account the three-dimensional (3D) regulation of genes through, for example, enhancers and silencers, which can be situated many kilobases away from a gene, or even on other chromosomes (Chang et al., 2025; van Arensbergen et al., 2014). Nevertheless, if this variant results in an increase in *SARM1* promoter activity, it could be very important for 16% of the population, based on an allele frequency of 0.08, as found in Project MinE DF1. While PMV13 does not appear to be associated with ALS, its high frequency makes it a useful variant for testing association with other neurodegenerative diseases.

The cells utilised in our study are cell lines that cannot recapitulate neurons *in vivo*. Even between the two cell lines we used, we found a stark difference in the *NMNAT2* promoter repressive activity, with clear repression in SH-SY5Y cells but none in HEK293T cells. Interestingly, it has been argued that HEK293T cells possess certain neuronal qualities (He & Soderlund, 2010; Shaw et al., 2002), so whether these or SH-SY5Y cells recapitulate *NMNAT2* or *SARM1* transcriptional regulation in neurons *in vivo* better is yet to be determined. Moreover, it has been proposed that the regulatory DNA sequences and transcription factors that interact with the *NMNAT2* promoter can vary depending on cell differentiation state (Chang et al., 2025). Thus, it is important to consider that transcriptional regulation of these two key genes may differ between cells and between neuron types. Indeed, it was recently demonstrated that ERK1/2 signalling can regulate *Nmnat2* transcriptional regulation in mouse dorsal root ganglia neurons but not cortical neurons, and that CREB mediates *Nmnat2* transcription in mouse cortical neurons but not dorsal root ganglia neurons (Yue et al., 2026). These authors suggest that this differential regulation may underlie the neurotoxic effects of MEK inhibitors used in cancer therapy on the PNS. Indeed, transcriptional regulation differences could underlie different neuronal subtype susceptibility more widely in neurodegenerative diseases, for example could ALS primarily affect upper and lower motor neurons, but not sensory neurons, because of different expression levels of *NMNAT2* and/or *SARM1*? We find the NMNAT2:NAMPT ratio is different in mouse dorsal root ganglia compared to superior cervical ganglia neurons, suggesting possible regional transcriptional differences (Antoniou et al., 2024). Another hypothesis is that differences between individuals in the transcriptional regulation machinery controlling *NMNAT2* or *SARM1* contribute to why many neuropathies present as a spectrum of motor and sensory phenotypes.

In summary, our study uses functional assays to assess important regulatory regions in the *NMNAT2* and *SARM1* promoters. We show that SNVs that occur naturally in the human population have the potential to substantially modify promoter activity and thus gene expression. Moving forwards, it would be valuable to confirm the effect of these mutations when integrated into the genome (for example using CRISPR-Cas9 or with cells from individuals who possess these specific variants). These non-coding region variants would then add to the body of coding region loss-of-function *NMNAT2* (Dingwall et al., 2022; Huppke et al., 2019; Lukacs et al., 2019) and gain-of-function *SARM1* mutations (Bloom et al., 2022; Gilley et al., 2021) associated with disease, and thus could help drive statistical power in future population genetics studies. With these two key players of PAxD harbouring much attention in the field of neurodegenerative disease as possible therapeutic targets, we also contribute data that might inform targeting them on a transcriptional level.

## Acknowledgements

LC, HM, JG, ERW and this work was funded by a Wellcome Trust Collaborative Award in Science (220906/Z/20/Z/WT) awarded to MPC, MMR, Ahmet Hoke and David Bennett. The authors would like to thank the Project MinE ALS Sequencing Consortium. We are grateful to members of the Coleman laboratory for many useful discussions contributing to this work, as well as to Saif Haddad for review of the manuscript.

MPC consults for Nura Bio, Outrun Therapeutics, Sironax and DRISHTI Discoveries, and the Coleman group is part funded by AstraZeneca and Bristol Myers Squibb for academic research projects; none of these activities relate to the study reported here.

---

**Supplementary Figure 1:**
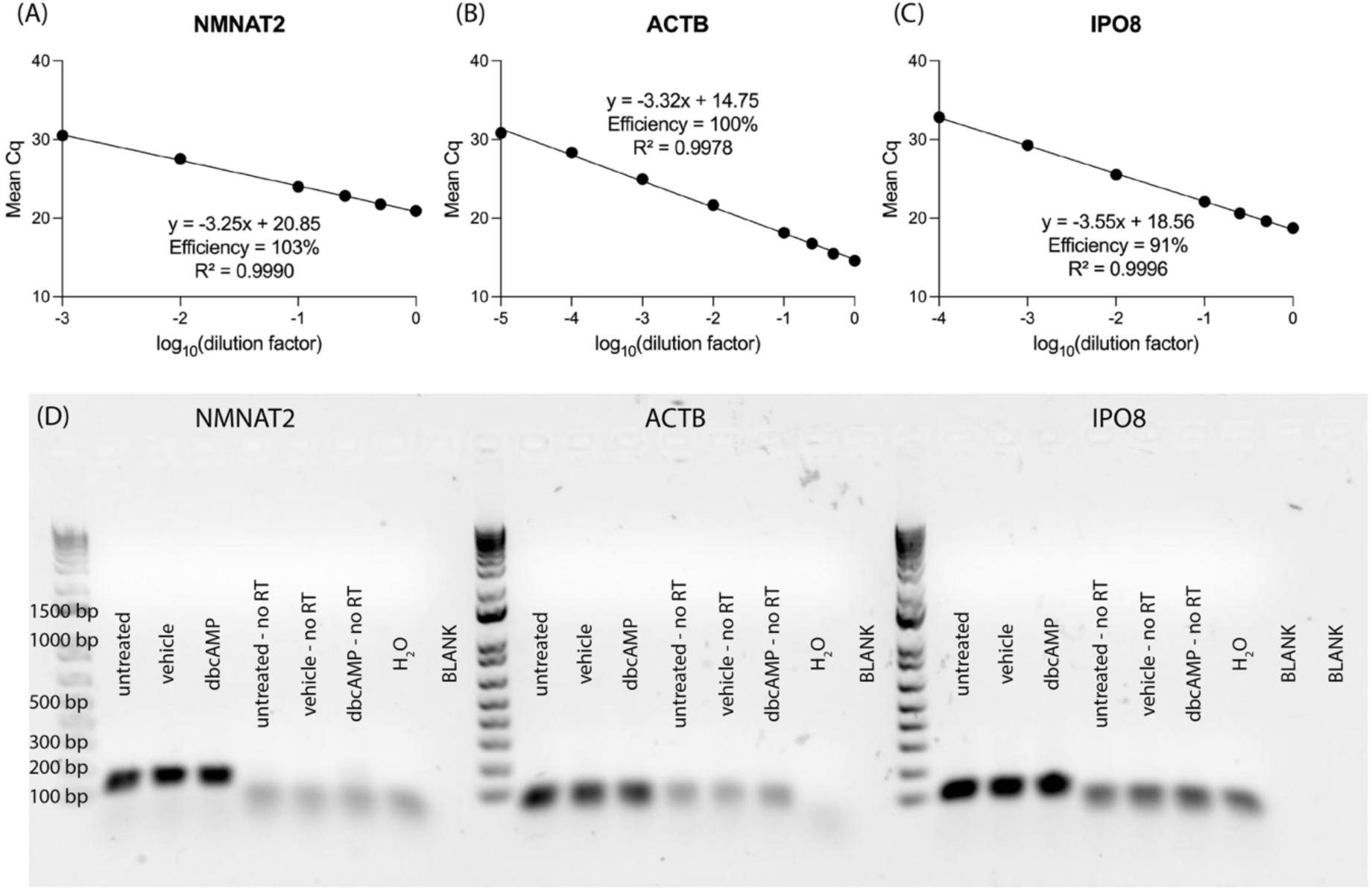
Quality controls for RT-qPCR. (A)-(C) cDNA from all the experimental groups of one replicate were pooled and subjected to serial dilution. Quantitative PCR was performed with each of the three primer pairs used in this study. Dilutions in which the standard deviation of technical replicates was ≥ 0.5 and/or multiple peaks were detected by melt curve analysis were removed from the plots as the concentration of the gene of interest was considered to be beyond the limit of quantification. For *NMNAT2* these were 0.0001X and 0.00001X, and for *IPO8* 0.00001X. Simple linear regression was performed in order to determine primer efficiencies. (D) Representative agarose gel images; RT-qPCR products were run on a 1% agarose gel in order to assess the number and size of PCR products. No RT = no reverse transcriptase control.

**Supplementary Figure 2:**
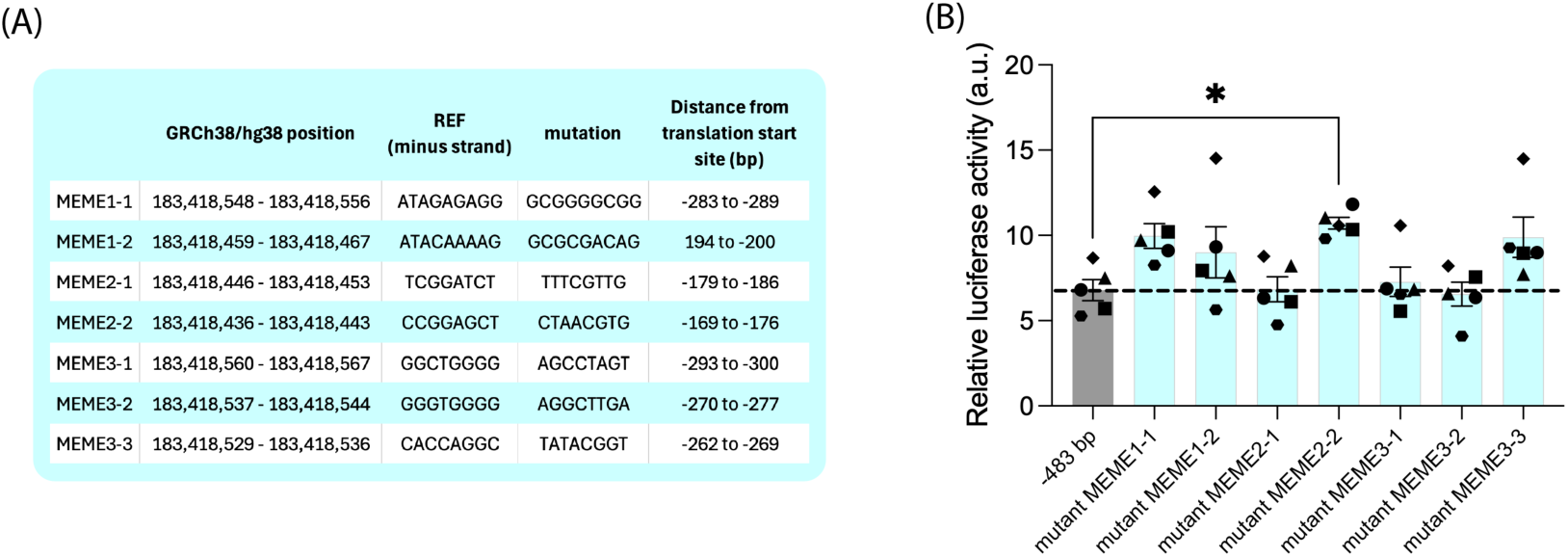
‘MEME’ sites, identified as potential transcription factor binding sites, do not independently contribute to maximal *NMNAT2* promoter activity. (A) Three prospective transcription factor binding motifs were identified (MEME1, MEME2, MEME3), found between -169 and -300 base pairs upstream of the ATG start codon. These sites were mutated to determine if they are important for *NMNAT2* promoter activity. REF = reference allele, mut = mutation, bp = base pairs. (B) Luciferase assays in SH-SY5Y cells suggest that mutations of MEME sequences mostly have no effect on *NMNAT2* promoter activity, except for MEME2-2, mutation of which results in a modest increase in activity. Data are presented as mean ± SEM. N = 5 (independent experiments are indicated by different shape data points). Statistical significance was determined by Kruskal-Wallis (*P* = 0.003). The statistical significance indicated on the graph (* *P* ≤ 0.05, ** *P* ≤ 0.01, *** *P* ≤ 0.001) was determined by Dunn’s multiple comparisons tests.

