## Supplemental Materials for "Human promoter analysis of the Programmed Axon Death genes *NMNAT2* and *SARM1*"

**NMNAT2 promoter sequence used in luciferase plasmids (5' to 3')**

**Variation from the reference genome (GRCh38/hg38 assembly) are in red and bold**

**5'**

GCCACTAATCTGCATATCATATTAGGGAAGATACAAACCCCATTATAAATATATATCAGTTCTTAG

**-2528 bp**

**1:183,420,795**

GAGTCTTCTATTTAAACTAGAAATTTTATGCGTATTAGGGTTGATTTGCTATTCATCATAATTTAACA

AATATTTACTCCACACCTACCACATGCTCTTGGATAAGAAAAAACACAGTGACCAACTCACTA

ATATGGGGTGGAAATTTATCCAAGAAACTTGAAGCCCAGGATAGGTACGTATTTGTACTTGTCA

ATCTCTAAGTTAAGAATTATTGGCTCTGGCTGGGTCTGTGGCCCATGCCTTTAATCTCAACAGT

TTGGGAGGCTGCCGTGGGTGGATTGCTTGAGGTCAGGAGTTCAAGACCAACCTGGGCAATA

TAGTGAGATCCCATCTCTACAAAAAACACGAAGAATTAGCTGGACATGGTGGCAAACACCTG

TAGTCCCAGCTACTTGCAAGGC**T**GAAGTGGGAGGATTGCTTGAGTCCAGGAGATCGAGGCT

GTGGTGAGCTGAGATCATGCCACTGCATTCCAGCCTGGGTGACAGAGTGAGATCCTGTCTCA

AAAAAAAAAAAAAAAAAAAAAAGAATTATCCTCTATATCCAATTCATCTGCAAAGAATTATTGGTT

**-1929 bp**

CTTTCTGTTGTATAAGCATAGAACATCTATCTTCCCCTACCATTCTCAATTTCAGTCTACTCATTG

TTTAAGCCAAATTTCAAAATCTATAAAACATATCCTGAGACCATCTACTTCTCGTTGCTCCCGTC

TCTATAAACCCTGGTCTCAGGACAACTCCACTTGCCTAGGCTACCAGATTTTCCTAAATGCTTT

CCTTGTCTTCCCATTTGCCCTCCCTTTCCACCCAGCAGCCAGAGTGATCCTATAAAAATGTAA

ATTACATATTATTCCCCTGCTTAAAAGCTTCAGTGGCTTCCTTTTGTTATTATTCCACTTAGAAAA

GTGTCTAACTCTTGCCCTAG**T**CTACAAGCCCCATTTGATCTGGCCCTGATCACCTGCTGCTCA

**-1555 bp**

**T missing only in -1555 bp construct**

GTCTTCTATCATTTTTCCCCTTGCCCATACTTTCTAGCATCACTGGCCTTTCATTTCTATGAATAC

AGTAAGCCCTTCCCTGTCCCTGAATACGTGAGCTAGCAACTACTCCTGTCGCTCAGGTCTCA

CTGAAATGTCACCTCCCCAGAGAGGCCTTCTCAGACTACTTGATTCGTCCTCTCTTCTCCACT

CACTCTCTGCTCTGTCAGAGAGTAAATTGCATTCACTTACTGTGCATTGCACAATTGCCATCTA

ATATTTATTGTCTATCTTCCTCCAGTAAAATGAAGACAGTGACTTTGCTTCACCAGCAGGGTTTA

TAGCAATACTTGACACATAATGAGCCCTCAAGAAACACTTGTTAAATGGTAAGTCTTTGTCTCAA

CTTTGGTAGCTTATGGAGGATCCCCATCATTTTCTTTTTTCTTTTTTTTTGTTTCGAGATGGATTCT
**-1116 bp**
CGCTGTCGCCCAGGTTGGAGTGCAGTGGTGCGATCTTCGCTCACTGCAACCTCTGCCTCC

CAGGTTCAAGCAATTCTCCTGCCTCAGCCTCCTGAGTAGCTGGGATTACAAGCACTCGCCA

CCATGCCCGGCTAGTTTTTGTATTTGCTTTTTAGTAGAGACAGGGTTTCACCGTGCTGGCCAG

GCTGGTCTCGAACTCCTGACCTCGTGATCCGCCCGCCTCGGCCTCCCAAAGTGCTGAGAT

TACAGGCATGAGCCACCGCACCCGGCGGATCCCCATAATTTTCTAATACAGTTTTAGTAACT

CTGCCCCCTTCATGTTCTCTCAAACGGATAGCATTGCTTCGCCCACGAAAAGGATCAGTATT

GCAGAGGAAGGGCTTAGATCACCCCTATCTCTCTGCAGCCAGCAGATCAAGATTTGGACTC

AACTCAATAGTTTGGAATCATCAGATAAAGGAAAAAGGAAAAGCAGAAATGAATACTACTGAAA

TAAACAATGGTCATTTCTCACTTAAAATTTTGAGGAAGAGAGAAAGAAAAAATACCTTAAGATCA

AACATTTTTTAACTTGGAGATGTTAGAGAAAGTAGGGGGGACAGAAAAATGTTAGAATGAGGA
**-480 bp**
GGAAAACTGAATCAAGATGAGGCAAGAGGCAGAAAGAGCTCACACCATGCTGTGGGCTCTT

GGTTCTGAGCTTGGATACCACGTCTTGCCTTCTGGATAAACTCTAAGGAAGACAGTGATGGAG

TGAAGTGGGCTGGGGGCGATAGAGAGGATGGGGTGGGGCACCAGGCGAGAGATGCGAA
**-286 bp**
GGAAGCCAGAACGAAAAGAGAGCGACCGAGGAGAGAAGAGAGCAGAGCAATACAAAAGC

AGCCTCGGATCTAGCCGGAGCTGCAAGCGTTAAGGGGAGGCGGAGAGTGACGCGGTTTG
**-170 bp**
CGTCTGGAGCGGCTCCTTGGAGTCCACAGCATCCACCGCCGGAGCCTCGCCTTCCTTTCT

CCCTCTGCAGACACAACGAGACACAAAAAGAGAGGCAACCCCTAGACCACCGCGAAGGA

CCCATCTGCACC**ATG** **3’**
**+1**
**translation start codon**
**1:183,418,267**

**<u>SARM1 promoter sequence used in luciferase plasmids (5' to 3')</u>**

**Variation from the reference genome (GRCh38/hg38 assembly) are in red and bold**

**5'**

CTTGAGGTCGGTGAAGGCGTCGAAGGGCTTCCCACTGCACAGCTCCTCCTCTGCTGGGGG

**-2501 bp**

**17:28,369,532**

CTGAGGTCTCCCTGGATGAAGGGTCTCAGGCCTTGAGTCTATCCCCTCAGGCTTAGAGGCG

CCCACCTCAGGCGCAGGGGCCTCTTCCTCAGGTTTCAGAACAGGTGTCTGCTCAGGATTC

CCTTTGGACTGGGCCTGGAGGTCAGAGGTCAGGGAGGGGCCCCCCACCTGTTCATGGAC

AGTGGCATTGTTTTTCTCCTCGCCATCGTCATAGACCGTGTACTCATCCTCCGGCATAGTGAA

CACATCCCCGCGAGTCACTGCAGAGAGTGGATGGTAGTGAGTCTCCAGCAGCAGGGGGC

ACCCCAGCCCACCCACCTGGGCTCTGAACACACCTTGGGGCTTGCACTCAGCCGTATAGT

CTGTGCAGCAGCTCTGGTAGTAAGAGCAGAGCTCGTCACACTGGCACTTCTTGTCCACGTTG

AAGCCCTCAGTGCAGCGGCCCTTGCATGACTCTATGAGGAAGGAGTGTCAGTCGGTGCCA

CCAAGCCCAGACCACCCTCGCCCTCCCTACATTGACCCAGATGGCCACCAACACTCCCC

TGTACCTTGGTCAGCCAGAGCAACCCATGCCAGCAGGGCCAGTATGAGAAGGGGTCTCAG

GGGTGCCATGGCAGGGCTTCTAGCTCAGTGCCTGGCAAGCTGGGCTCTGGTCTCCCTGAA

**-1823 bp**

GTCTCCGCTCTGATGCCTGAGGAAGGGAGGGAGAGGCAGAGACAGGGAAGGAGGGCACT

GGAGAAGAGGAACTGCCTTTTCTGCTGCTCTGTTTGCTCAACCTCCAGCCCATCCCCTCTG

CCCCCTCCAGCGGCTGCTGCAGCAAAGGTCACATTCCTGGAACACTGGGCCTGGGCGAG

CTGGGAGATAAGACCTTTGCCAAGCTCAGAATCATTAGGTCATCGGAAGGGGAATTAGCACC

GTGGATCTGGAGGGCAGAAAGAAGGCTCATTGGGCAACAGCTTTATCTCTCTGAGTCTTAGTT

CATTTTCAGTAAAATAGGATTAACAAAGCCTTGCTTCATGGGGTGTTGGGAGATTACATGAACT

GGACCAGACAAAATGCCCAGTAGCTGGAGCAGCCACAAATTCCTTCTCCAAACATAGCTATT

GATCCATGATTGTTTGATCAGACACATCCCTGGCTGGGTGCTCATTCGCCATTATTTATTTAGG

GCAGGAAAAAGGGAGTGGGAGGAGAGATTGTAAGCACTCTGGGGAATTTTATTTTTTAGCATA

AAAGAACAAAGTTTCATTTCTGGGTTCTTCTCTTGGGTGCATCAATCTCCCCAGGCTGGAAATT

TGGGTAAATATCAACCCACTGGACTGGATTTAATTTCTGAGTCTTTTCTGAGCAGGCCACTCC

CTGTCCCCAGGCTCAGTCTCCCCATCTGTAAAGAGTGGGCTTGTTCAACACACTTTTAGACT
**-1093 bp**
CTCGAGAGAAATATGATCCCTCTACCTGGAAAATGCCCACTGGCATGAAACACCTGATTTTGT

GCCCAATTTCAGGTGTCCACAGAACTTCTGAAATCCTCCCATCACCTGAAAAAAGTGATCCT

GAGATTTTAGCTGACTTTTTCTGTTCTGTAATCAGAGGGGACTGTGATTTGGGCCAATTTCTTCC

TCATGGGCTTCAGTGGAGTCAGCTGTGGAATGGAACATCAATTCCCACCCCATAGGCATGTT

GTGAGGTTTCGCTAAGAAACAGCCATAAAAGGGCCTTAAGAGTTATGATGATTGTTTTAACCAT

TTAGTAGGGAGGGGGCATCCAGCGGTGTTGCTCCCACGTACCGATGTCACGTGGGGCGG

GGAGGCGGGGCGGAGAACGGAGAGCGTCCTCTCATTCTCCACCCCCTTCCTCCAAGTCC

AGCGCAGGGGGAGGTGGCTGCGACTGACTGAGGGGCTAGGGAAGGCTGGCCAGGGGC

GTGGGCGTGGCTGGGGACGGCTTGGGGGTGGGGCTGCGCAGGAGGCTGGAGGAGCCC
**-531 bp**
CGCGGGACCCAGAGGCGGGGCGTCGGCCCGGGGACC**G**CTGCTCCTCCGGGGCGTGG
**C**
CTGCAGGAGGCTGGAGGAGCCCCGCGGGACCCAGAGGCGGGGCGTCGG˄CCGGGGAC
**extra C compared to reference genome**

CGCTGCTCCTCCGGGGCGTGGCTGCTGCCGAGCATCTCCCAGCTCAGCCGAGCCCGTG

CCCAGGCCACGCTTTGTTCCAGCCGCCGCCTCCTCTACCCTACGGCGTCCGGAGCCATC
**-286 bp**
CCTCGCCTGCTCGCTCTCTCCTTTCGCCCACTCCCTGCATCTGGGCCTGCATCACCTTTGC

CAACCGCTCCCCCGATCCTGCCGACACTCCTCCCCCAAACTTCTGACCGGCACCCTTGC

CTGGTACCCTTCTCTCCATTCCTCCCCCTCCATCTTCTTTCCCCGACCCCTCTCGGGTCCCT

CTTTTCCCAAAACCCGGGTCTCTCCGCGTGGCCCCGCCTCCAGGCCGGGGATGTCCCCC

GCGGCCCCGCGCCC**ATG** **3’**
**+1**
**translation start codon**
**17:28,372,033**
